# Haplotype-resolved diploid genome inference on pangenome graphs

**DOI:** 10.1101/2025.11.26.690754

**Authors:** Ghanshyam Chandra, William T. Doan, Daniel Gibney

## Abstract

Recent algorithmic advancements have shown how to utilize pangenome graphs in combination with the haplotype reconstruction framework of Li and Stephens to accurately reconstruct a haplotype from a reference pangenome graph and a set of input reads. However, significant work remains in developing techniques that utilize a pangenome graph to obtain a pair of phased haplotypes called a diploid pair.

**Results:** We introduce new problem formulations and scalable algorithms for inferring phased diploid genomes from a pangenome graph and a set of input reads. We implement them in our tool `DipGenie`. The key idea is to jointly optimize genotyping and phasing along global paths through the pangenome graph, guided by a biologically motivated recombination budget that constrains inferred haplotypes to plausible mosaics of reference haplotypes. We evaluate `DipGenie` on real Illumina short-read data from the highly polymorphic MHC region in 22 leave-one-out diploid experiments, benchmarking against three tools that also operate on graph structures: `VG`, which samples haplotypes directly from the pangenome graph, and `PanGenie + Beagle` and `Paragraph + Beagle`, which derive local graphs from a VCF panel for per-site genotyping and delegate phasing to a statistical method. At full coverage, `DipGenie` achieves a geometric mean switch error rate (SER) of 0.86%, which is 5.7*×* lower than `PanGenie + Beagle` (4.88%), 7.9*×* lower than `VG` (6.77%), and 13.2*×* lower than `Paragraph + Beagle` (11.35%). For structural variant calling, `DipGenie` leads with a geometric mean F1-score of 0.571, compared to 0.470 (`PanGenie + Beagle`), 0.450 (`VG`), and 0.379 (`Paragraph + Beagle`). These advantages hold at every coverage level tested.

**Availability and Implementation:** https://github.com/gsc74/DipGenie.

## 1 Introduction

Pangenome graphs provide a useful and space-efficient representation of haplotype-resolved genomic references. They can accurately capture genetic variation, including SNPs, indels, and structural variants, while also leveraging similarity between references to reduce the required storage space [37]. Genome graphs further enable the alignment of reads to all references simultaneously in a single sequence-to-graph alignment procedure. These properties have been used extensively in population-scale variant analysis and reference representation studies [12,20,50], and have been developed into popular tools [6,8,9,17,25,27–29,32,35,46,51].

Genotyping is a method for identifying unknown variants at reference loci within a given sampled input [39]. It can be performed using either alignment-free methods (e.g., k-mer counting) or alignment-based methods, where it may use haplotype reference panels or a pangenome graph. Genotyping is an important problem in computational biology, as it enables downstream applications such as disease risk analysis and ancestry inference. Several large efforts have been initiated to create reference panels to improve genotyping quality, including the International HapMap Project, the 1000 Genomes Project, the Haplotype Reference Consortium (HRC), UK10K, GenomeAsia100K, the Human Pangenome Reference Consortium (HPRC), the Human Genome Structural Variation Consortium (HGSVC), and the Genome India Project [1, 2, 13, 26, 37, 38, 55, 56]. Genotyping methods have been widely implemented in software [4, 5, 18, 19, 27, 28, 31, 40, 42, 51].

Recently, pangenome graph-based genome inference methods have been shown to improve genotyping accuracy, particularly for structural variant detection in short-read datasets [18, 19, 27]. Chandra et al. [8] proposed a combinatorial optimization-based formulation of genotyping as a haplotype inference problem that simultaneously seeks to explain the given read set and limit the number of recombinations used. They further provide an NP-hardness proof and integer linear as well as quadratic programming formulations.

A further extension to genotyping is phasing, which refers to assigning variants to their respective haplotypes. For humans, that means finding which haplotype in a diploid pair a given variant occurs in. In the traditional approach, a large set of accurately phased haplotypes is used to guide phasing, as well as to infer unobserved variants (imputation) [4, 5, 22, 30, 31, 40, 49]. There is currently limited work on using pangenome graphs for phased genotyping that improves accuracy over traditional approaches. For instance, the approach of Chandra et al. [8] is limited to inferring only haploid solutions. While they briefly discuss extensions to polyploid genomes, their formulation does not incorporate phasing information, which restricts its ability to infer fully phased diploid genomes. Siren et al. [52] proposed a problem formulation and algorithms for inferring phased polyploid genomes from a pangenome graph; however, their formulation accounts only for block-wise phasing with a block size of 10 kbp. This motivates the need for new problem formulations and algorithms that enable diploid genome inference while accounting for end-to-end phasing information.

We address this gap by formulating diploid genome inference as a combinatorial optimization problem on pangenome graphs that accounts for end-to-end phasing. We implement the proposed algorithms in a `C++` tool, `DipGenie`, and evaluate it against three competitive tools that also operate on graph structures: `VG` [52], which samples haplotypes directly from the pangenome graph, `PanGenie + Beagle` [4, 18], and `Paragraph + Beagle` [4, 10], both of which derive local graphs from a VCF panel for per-site genotyping and delegate phasing to a statistical method. We highlight our contributions as follows:

– We propose a novel problem formulation for diploid genome inference, design a corresponding algorithm, and provide a theoretical framework to compute *a posteriori* approximation ratios that certify solution quality after each run.
– We conduct a comprehensive evaluation in the MHC region, the most polymorphic region of the human genome [16], performing leave-one-out experiments across 22 diploid samples with real Illumina reads at coverages of 1 ×, 2×, and total (2.8 to 12.4 ×). We benchmark phasing and structural variant calling accuracy against the three tools above.
– We show that `DipGenie` outperforms all competitors at every coverage level. At full coverage, it achieves a geometric mean switch error rate of 0.86%, up to 5.7 × lower than `PanGenie + Beagle` (4.88%), 7.9 × lower than `VG` (6.77%), and 13.2 × lower than `Paragraph + Beagle` (11.35%). For structural variant calling, `DipGenie` leads with a geometric mean F1-score of 0.504–0.571 across coverages, compared to 0.318–0.470 (`PanGenie + Beagle`), 0.319–0.450 (`VG`), and 0.275–0.379 (`Paragraph + Beagle`). These gains arise because `DipGenie` jointly optimizes genotyping and phasing along global graph paths under a biologically motivated recombination budget, whereas `VG` phases only within local 10 kbp blocks and the genotyping pipelines impose no graph-structural phasing constraint. `DipGenie` is also the only method whose mismatch rate improves monotonically with sequencing depth.

### 1.1 Problem formulation

Our tool utilizes the Li and Stephens population genetics model [36] which infers unknown haplotypes as mosaics of reference haplotypes. In this model, a switch in the inferred haplotype from one reference haplotype to another corresponds to a recombination event.

We consider an acyclic directed graph (DAG) representation of a pangenome reference. Let *G* = (*V, E*) be a DAG. Each vertex is labeled over an alphabet *Σ* by a function *σ* : *V* → *Σ*^∗^. Note that cyclic regions of more general graphs can be unrolled using standard techniques, with some additional overhead. We use ℋ to denote a given set of paths in *G*, referred to as *reference haplotypes*. Together, these define a haplotype-resolved pangenome DAG *G* = (*V, E, σ*, ℋ). We assume that every vertex *v* ∈ *V* is contained in at least one path in ℋ, and that every edge (*u, v*) ∈ *E* connects two consecutive vertices along some path in ℋ.

For an arbitrary path *P* = *u*_1_, *u*_2_, …, *u*_*k*_ in *G*, we overload the notation so that *σ* denotes the string spelled by *P* ; that is, *σ*(*P*) = *σ*(*u*_1_)*σ*(*u*_2_) … *σ*(*u*_*k*_). By our stated assumptions, we can associate a haplotype path with every vertex in *P* ; that is, *u*_1_.*h*_1_, *u*_2_.*h*_2_, …, *u*_*k*_.*h*_*k*_, where *h*_1_, *h*_2_, …, *h*_*k*_ ∈ ℋ. Here, the notation *u*_*i*_.*h*_*i*_ indicates that vertex *u*_*i*_ is traversed as part of the haplotype path *h*_*i*_. We use *r*(*P*) to denote the number of *recombinations*, that is, haplotype switches, on *P* :

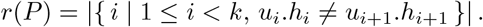

#### Haploid path inference

We summarize the formulation for haplotype path inference introduced by Chandra et al. [8]. In that work, one is given a haplotype-resolved pangenome DAG *G* = (*V, E, σ*, ℋ), a set of target strings 𝒮, and a non-negative integer *c* indicating a recombination penalty, and seeks to output an inferred path *P* such that

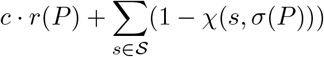

is minimized, where *χ*(*s, σ*(*P*)) = 1 if *s* occurs as a substring of *σ*(*P*) and 0 otherwise. The value *c* is picked empirically to produce a reasonable number of recombinations.

#### Diploid path inference

The problem of finding a diploid pair of sequences naturally generalizes, in some aspects, the task of inferring a single haplotype path. Rather than identifying one path *P*_1_, we now seek two paths, *P*_1_ and *P*_2_. However, a challenge arises if the objective is simply to maximize the number of target strings captured by either *P*_1_ or *P*_2_. For example, if the majority of target strings are captured by *P*_1_, then *P*_2_ is largely unconstrained and may follow any path in *G* that satisfies the recombination limit. Some mechanism is therefore required to ensure a reasonable degree of similarity between the two paths.

To address this, we consider the target strings as partitioned into two categories: *homozygous* target strings, which should occur in both *σ*(*P*_1_) and *σ*(*P*_2_), and *heterozygous* target strings, which should occur exclusively in one of *σ*(*P*_1_) or *σ*(*P*_2_). We later describe the method for performing this partitioning.

We formulate the diploid path inference problem as follows.

*Problem 1 (Diploid path inference)*. Given a haplotype-resolved pangenome DAG *G* = (*V, E, σ*, ℋ), two disjoint sets of target strings 𝒮_hom_ and 𝒮_het_, and a non-negative integer *R* indicating the maximum total number of recombinations, output two inferred paths *P*_1_ and *P*_2_ such that

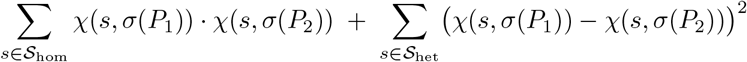

is maximized and

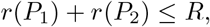

where *χ*(*s, σ*(*P*)) = 1 if *s* occurs as a substring of *σ*(*P*) and 0 otherwise.

This formulation balances coverage and redundancy between the two inferred paths, ensuring that both contribute meaningfully to representing the diploid sequence. The first term rewards shared coverage of homozygous target strings, ensuring consistency between the two inferred haplotypes, while the second term encourages complementary coverage of heterozygous targets, promoting diversity between the paths. See Figure 1 for an example. We further eliminate the empirical tuning of the parameter *c* in [8] by using a better motivated constraint, as described in the section on selecting the correct number of recombinations.

**Fig. 1.**
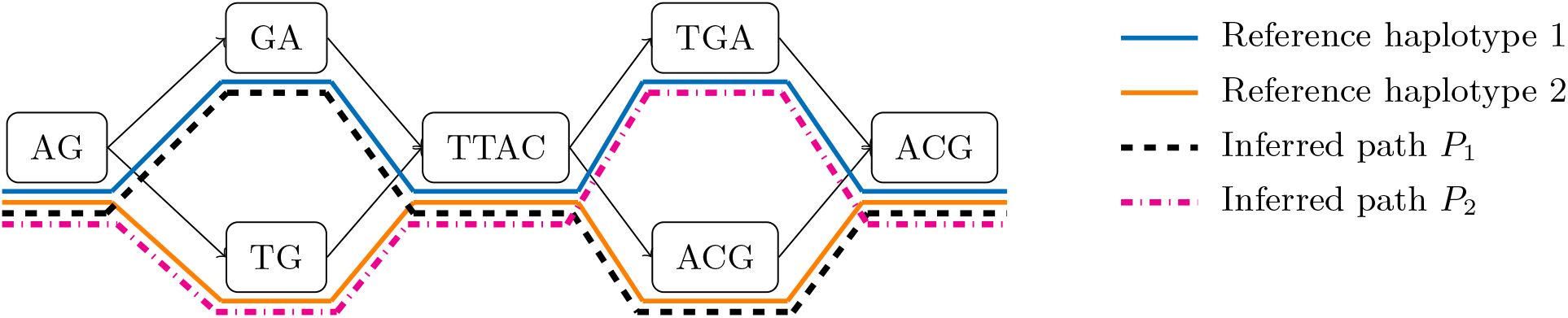
In the example above, the indicated solution uses two recombinations, one in *P*_1_ and one in *P*_2_. We have *σ*(*P*_1_) = AGGATTACACGACG and *σ*(*P*_2_) = AGTGTTACTGAACG. If 𝒮_hom_ = *{*GA, GAC*}* and 𝒮_het_ = *{*TAC, TGT*}*, then GA contributes to the objective value, since it appears as a substring in both *σ*(*P*_1_) and *σ*(*P*_2_). The string GAC does not contribute, as it occurs only in *σ*(*P*_1_). The string TAC does not contribute because it occurs in both *σ*(*P*_1_) and *σ*(*P*_2_), while TGT contributes since it appears only in *σ*(*P*_2_). Hence, the total objective value of this solution is two.

## 2 Methods

This section outlines an algorithmic approach to Problem 1. The NP-hardness of Problem 1 follows through a straightforward reduction from the haplotype path inference problem defined by Chandra et al. [8]. An exact integer quadratic programming solution (IQP) is provided in this work’s supplementary material. Unfortunately, this exact solution fails to scale well to larger instances. We therefore focus on a more scalable approach.

### 2.1 Expanded graph

We briefly outline the construction of the *expanded graph G*_*E*_ = (*V*_*E*_, *E*_*E*_) and the *anchor set*. We refer the reader to [8] for more details. The main idea is to decompress the given haplotype-resolved pangenome DAG *G* = (*V, E, σ*, ℋ) by explicitly representing each haplotype path *h* ∈ ℋ as an independent chain of edges with weight 0. A single source vertex *source* is connected to the start of each such chain, and a single sink vertex *sink* is connected from the end of each chain. Next, weight 1 edges, referred to as *recombination edges*, are added between these haplotype paths to represent possible recombination events between corresponding loci.

For every target string *s* ∈ 𝒮, all of its occurrences within any haplotype path are represented as subpaths. Collectively, these subpaths form the *anchor set*. We use hit(*s*) to denote the set of subpaths corresponding to occurrences of *s* ∈ 𝒮. Following [8], we restrict our model to hits that are fully contained within individual haplotype paths in ℋ. In our experiments, we use (*w, k*)-minimizers shared between the reference haplotypes and the input reads as the target string set 𝒮.

### 2.2 K-mer partitioning

Our *k*-mer partitioning approach begins by constructing a histogram in which the *i*-th bucket records the number of *k*-mers in 𝒮 that appear exactly *i* times in the read set. To classify the *k*-mers, we fit a mixture model containing:

1. a normal distribution with mean *µ*, corresponding to heterozygous *k*-mers, and
2. a normal distribution with mean 2*µ*, corresponding to homozygous *k*-mers.

After fitting the model, the contribution of each component distribution to every histogram bin is computed, allowing *k*-mers to be classified as heterozygous or homozygous based on their frequency. We refer the reader to the work of Chikhi and Medvedev [11] for similar techniques used for *k*-mer classification. For lower coverage, we find empirically that the simpler technique of treating the first histogram bucket, corresponding to *k*-mers with a single occurrence, as containing heterozygous *k*-mers and the remaining buckets as homozygous works well in practice.

### 2.3 Diploid color collector

We next introduce a very similar problem to Problem 1 that is more amenable to dynamic programming techniques. The key advantage is that a dynamic programming solution no longer needs to keep track of multiple partially matched subpaths, thereby simplifying both the state representation and the computation.

*Problem 2 (Diploid Color Collector Problem)*. Given as input:

– Disjoint color sets *hom* and *het* with *hom* ∪ *het* = {1, 2, …, *C*} where *C* is the total number of colors;
– A recombination limit *R* ≥ 0;
– A DAG *G* = (*V, E, w, c*) with edge weight function *w* : *E* → {0, 1} and vertex color assignment *c* : *V* → 𝒫({1, 2, …, *C*}), i.e., *c* maps a vertex to a subset of {1, 2, …, *C*};

find a pair of paths *P* = (*P*_1_, *P*_2_) in *G* such that the objective function

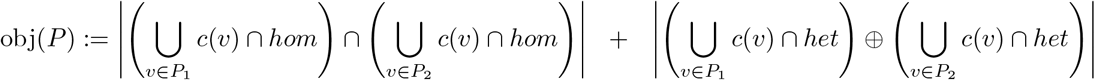

is maximized.and

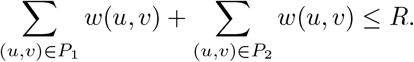

We transform our instance of Problem 1 to an instance of Problem 2 by beginning with the expanded graph representation described in the section on problem formulations. We start by assigning a unique integer identifier (henceforth the “color”) *γ* to every *s* ∈ 𝒮. Thus, *C* = |𝒮|. We use the classification of strings in 𝒮 into 𝒮_hom_ and 𝒮_het_, to define color sets *hom* and *het*. Next, we apply the following transformations (see Figure 2).

**Fig. 2.**
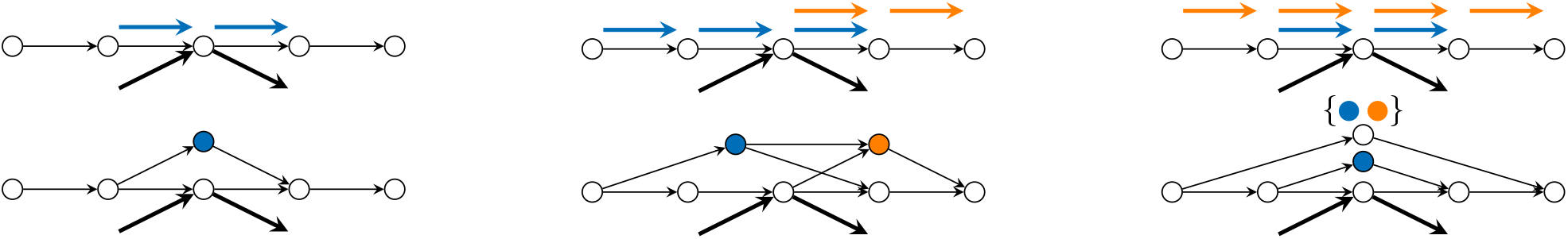
Graph transformations used to reduce the string-based Diploid Path Inference Problem to the Diploid Color Collector Problem. The top panel illustrates a target string occurrence (a “hit”) mapping to a sequence of nodes in the expanded graph. The bottom panel demonstrates how this sequence is compressed and transformed into a single vertex assigned a specific “color” corresponding to that target string. By applying this transformation to all target string hits and propagating these colors across overlapping regions, complex string-matching constraints are successfully translated into localized vertex-color matching constraints.

**Fig. 3.**
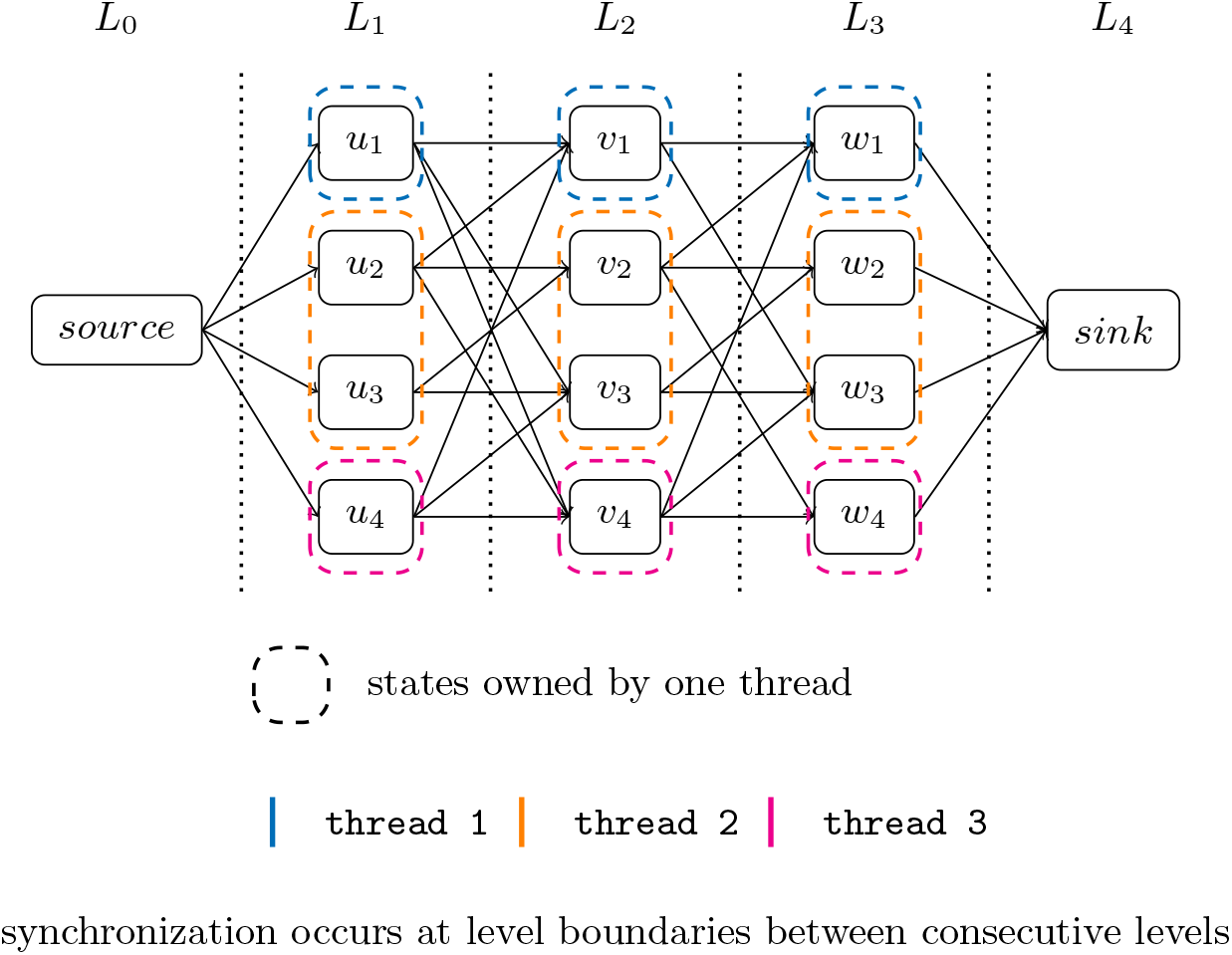
Illustration of the leveled DAG structure used in the dynamic programming algorithm and the corresponding parallel execution strategy. Vertices are grouped into levels *L*_0_, *L*_1_, …, *L*_4_, and transitions are processed only between consecutive levels. Within a level, dynamic programming states are partitioned among threads according to vertex ownership, shown by dashed boxes. A thread may own multiple vertices in the same level. Threads update their assigned states in parallel and synchronize only at level boundaries before advancing to the next level.

– For every *s* ∈ 𝒮, and for every path *u*_1_, *u*_2_, …, *u*_*k*_ ∈ hit(*s*), we add a new vertex *v* assigned the color corresponding to *s*, and add edges (*u*_1_, *v*) and (*v, u*_*k*_). We refer to these added paths as *bridges*.
– We then scan each haplotype path *h* in *G*_*E*_ in topological order. If two bridges (*u*_1_, *v*), (*v, u*_*k*_) and (*w*_1_, *v*^′^), (*v*^′^, *w*_*s*_) created for *h* overlap, that is, *w*_1_ *< u*_*k*_ *< w*_*s*_, we add an edge (*v, v*^′^).
– Next, we again scan each haplotype path *h* in *G*_*E*_ in topological order. If one bridge (*u*_1_, *v*), (*v, u*_*k*_) created for *h* is fully contained within another bridge (*w*_1_, *v*^′^), (*v*^′^, *w*_*s*_), that is, *w*_1_ ≤ *u*_1_ and *u*_*k*_ ≤ *w*_*s*_, we augment *c*(*v*^′^) by adding all colors contained in *c*(*v*). This propagation is repeated until all color sets stabilize.

#### Our approach to Problem 2

We next describe our dynamic programming approach for Problem 2. The key idea is to traverse two paths through a leveled DAG under a shared recombination budget, maximizing homozygous colors that co-occur and heterozygous colors that appear in exactly one path. This approach treats colors in different levels as distinct, which will lead to approxaimte rather than exact solutions.

##### 1. Leveling the DAG

Starting from the DAG constructed in the haploid case, we form a strictly leveled DAG by inserting dummy vertices as needed. We denote the resulting graph as

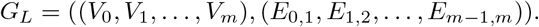

Level *V*_0_ contains the single source vertex, and each *E*_*i*−1,*i*_ contains all edges connecting *V*_*i*−1_ to *V*_*i*_.

##### 2. Dynamic programming

Let *hom*(*v*) and *het*(*v*) denote the sets of homozygous and heterozygous colors associated with vertex *v*, respectively. Let *D*_*l*_[*v*_1_, *v*_2_, *r*] denote the maximum objective value achievable by a pair of paths ending at vertices *v*_1_, *v*_2_ ∈ *V*_*l*_ using at most *r* recombinations in total. Initialize *D*_0_[*source, source, r*] = |*hom*(*s*)| for all 0 ≤ *r* ≤ *R*. For each level *l >* 0, all pairs of vertices *v*_1_, *v*_2_ ∈ *V*_*l*_, and all 0 ≤ *r* ≤ *R*, set

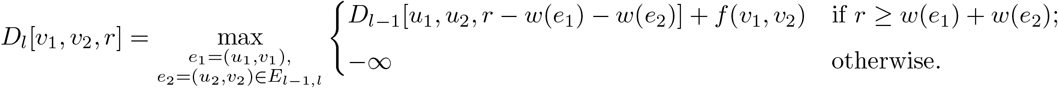

where

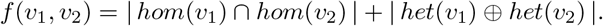

##### 3. Path recovery

For each recombination budget *r* ∈ { 0, …, *R* }, recover the corresponding pair of paths from *D*[*sink, sink, r*] and compute its true objective value. The pair with the best objective value is returned as the final solution.

The dynamic program above considers all edge pairs between consecutive levels, resulting in a total running time of

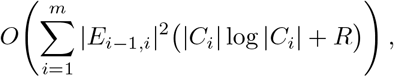

where 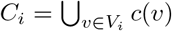. By maintaining only the weighted recombination edges for each path and updating one level at a time, the space complexity is *O*(max_1≤*i*≤*m*_(|*V*_*i*_|^2^ · *R*^2^ + |*C*_*i*_|)).

#### Diploid *a posteriori* approximation certificate

We provide an *a posteriori* bound quantifying how the multiplicity of color occurrences affects the quality of the diploid solution. Let *X* = (*X*_1_, *X*_2_) where *X*_1_ and *X*_2_ are two arbitrary paths in *G*_*L*_. Let occ_*γ*_(*X*_*i*_) denote the number occurrences of color *γ* on path *X*_*i*_ for *i* ∈ {1, 2}. We use *hom*(*X*) to refer to the subset of homozygous colors appearing in *X*_1_ or *X*_2_ and *het*(*X*) to refer to the subset of heterozygous colors appearing *X*_1_ or *X*_2_. We first define the following quantities:

#### Homozygous quantities

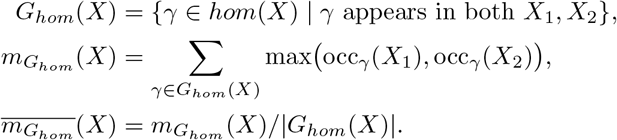

#### Heterozygous quantities

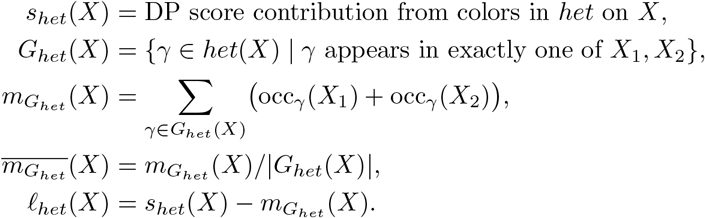

With these values defined, our *a posteriori* approximation ratio, obtained after finding a solution, is summarized in Theorem 1. Here, *a posteriori* means that after obtaining a solution we can evaluate how close it is to optimal, despite not knowing the optimal solution. This differs from standard *a priori* approximation ratio guarantees that upper bound the approximation ratio prior to obtaining a solution.

Let *P* = (*P*_1_, *P*_2_) denote the pair of paths returned by the approximation algorithm for Problem 2, and let 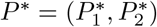 denote an optimal pair.

##### Theorem 1.

*Let P be a solution path pair obtained by our algorithm for Problem 2, and let P* ^∗^ *be an optimal solution. Let* 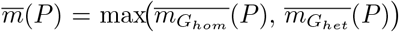 *be the maximum average multiplicity of contributing homozygous or heterozygous colors on P, and let ℓ*_*het*_(*P*) *denote the heterozygous score loss. Then*

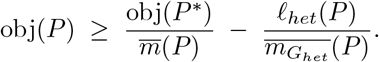

A formal proof is provided in the supplementary material. We report the a posteriori approximation certificates achieved by our approach experimentally in the results section.

### 2.4 Selecting the correct number of recombinations

Although taking the value of *r* ∈ { 0, …, *R* } that maximizes the objective is optimal for Problem 2, our ultimate goal is accurate diploid path recovery. To avoid overfitting and to reflect realistic recombination patterns, we estimate the expected number of recombinations using the Li and Stephens population HMM model [36]. Assuming a uniform per-base recombination rate *r* and effective population size *N*_*e*_, the expected number of recombinations over a reference of length *L* with |ℋ| donor haplotypes is

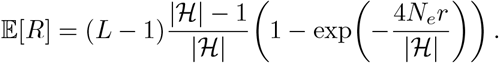

Using parameters consistent with prior studies (*L* = 4.9 Mbp, = 47, |ℋ|, *N*_*e*_ = 5 × 10^3^, *r* = 4.6 × 10^−9^ Morgans/bp), this yields 𝔼[*R*] ≈ 9. For diploid path inference, we use 2𝔼[*R*] ≈ 18. A full derivation and parameter justification are provided in the supplementary material.

### 2.5 Parallelization strategy

To improve scalability on large pangenome graphs and high-coverage read sets, we parallelize several stages of the `DipGenie` pipeline using shared-memory parallelism with OpenMP. During anchor generation and classification, the computation over minimizers and reads is distributed across threads, where each thread independently processes subsets of minimizers and stores intermediate results in thread-local buffers before merging them into the final data structures. This avoids excessive synchronization overhead while enabling efficient parallel processing of large collections of reads and haplotypes. We additionally parallelize the construction of anchor mappings and read-support structures, which represent a significant portion of the preprocessing cost on large datasets.

Our diploid dynamic programming algorithm is parallelized across states within each graph level of the expanded graph representation. The algorithm processes vertices in topological order and uses rolling dynamic programming buffers to reduce memory usage. Transitions from each state are evaluated in parallel, while synchronization is handled through striped locking to safely update shared dynamic programming states with low contention. To further improve scalability, each thread maintains local edge pools for traceback information, avoiding repeated synchronized memory allocations during dynamic programming updates. Since dependencies occur only between consecutive graph levels, synchronization is required only at level boundaries, allowing substantial parallel speedups while preserving correctness of the dynamic programming recurrence.

## 3 Results

We implemented our diploid path inference algorithm in C++ as `DipGenie`. The tool accepts a pangenome reference in GFA format together with short reads from a diploid sample, and outputs two inferred haplotype sequences in FASTA format. Target strings are identified using (*w, k*)-window minimizers [47] (*w*=25, *k*=31), and the recombination budget is *R*=18 (see Expected number of recombinations). All experiments were performed using 128 threads on a single-socket 64-core Intel Xeon Platinum 8592+ processor with 1 TB RAM. Unless otherwise noted, all summary statistics are geometric means (GM) across 22 samples.

### 3.1 Datasets and experimental design

The major histocompatibility complex (MHC) is among the most polymorphic regions of the human genome [16], presenting a challenging test bed for haplotype inference. We obtained MHC haplotype assemblies for 24 diploid samples [34] along with whole-genome Illumina reads for 22 of these from the HPRC year-2 data release (Supplementary Table A1). For each sample, we constructed a leave-one-out pangenome graph from the remaining 23 samples and the CHM13 reference [43]; correspondingly, we used `vg deconstruct` [28] to project graph variants onto the linear reference and `vcfbub` [24] to decompose the resulting nested variant sites. The graph serves as the reference panel for graph-based tools, and the decomposed VCF serves as the linear reference panel for genotyping-based tools. We derived ground-truth VCFs by aligning each held-out sample’s haplotype assemblies to the CHM13 reference with the Cactus pipeline [29]. The downloaded reads for 22 samples were mapped to `GRCh38.p14` using `minimap2` [35]; paired-end reads aligning to the MHC region were extracted and downsampled to 1×, 2×, and total (full) coverage. For accurate phasing of genotyped variants, we extracted the genetic map for the MHC region on `chr6` from the CHM13 recombination map [3]. Graph and read statistics appear in Supplementary Table A1, and all datasets are available on Zenodo [7].

We benchmarked `DipGenie` against three methods that also operate on graph structures. The first approach is graph-based haplotype sampling: a) `VG` [52] divides the pangenome graph into non-overlapping blocks of target length 10 kbp, identifies graph-unique *k*-mers [33] within each block, and counts their occurrences in the reads to classify each haplotype as absent, heterozygous, or homozygous; the highest-scoring haplotypes are greedily selected with scores iteratively adjusted to maintain balanced allele representation, and the sampled local haplotypes are recombined across blocks to construct two full personalized paths; test VCFs for `VG` are generated via the Cactus pipeline, identically to the ground truth. The remaining two tools derive local graphs from a VCF panel for per-site genotyping and delegate phasing to a statistical method: b) `Paragraph + Beagle`: reads are aligned to the CHM13 reference using `BWA-MEM2` [54] and SNP likelihoods are called with `bcftools` [14]; `Paragraph` [10] then constructs a directed acyclic graph around each structural variant’s breakpoints from the VCF panel, aligns reads to this graph, and counts supporting reads along reference and alternate paths to compute SV genotype likelihoods; c) `PanGenie + Beagle`: `PanGenie` [18] identifies *k*-mers unique to each panel haplotype path and uses their *k*-mer counts in a Hidden Markov Model (HMM) to compute genotype likelihoods at each variant site. Both tools operate on the decomposed linear VCF panel derived by `vg deconstruct` and `vcfbub` [24, 28], and both produce unphased genotypes that are subsequently phased with `Beagle` v5.4 [4, 5] using the MHC genetic map [3].

Together, a), b), and c) cover the major approaches for pangenome-based diploid inference: graph-based haplotype sampling directly on the pangenome graph, and per-site genotyping from local graphs derived from a VCF panel with separate statistical phasing. We excluded `Locityper` [45], a targeted genotyper designed for individual polymorphic genes, because it evaluates all pairs of existing panel haplotypes and outputs the best-matching one based on sequencing errors, insert sizes, and read depth; applied to the MHC, it would return an unmodified pair from the reference panel rather than recombined haplotypes tailored to the sequenced individual.

### 3.2 Evaluation metrics

We evaluated the accuracy of inferred haplotypes with three metrics. We used `whatshap compare` [41] to assess phasing quality over the set of heterozygous variants common to truth and the inferred VCFs. The tool partitions common phased variants into non-singleton intersection blocks, defined as maximal runs of at least two heterozygous variants that are phased in both files. a) The switch error rate (SER, %) is the number of switch errors divided by the number of phased pairs of variants assessed; a switch error occurs between two consecutive heterozygous variants in an intersection block when the phase relationship between them in the inferred haplotypes disagrees with the truth. b) The mismatch rate is the block-wise Hamming distance expressed as a percentage: the number of heterozygous positions within intersection blocks at which the inferred allele-to-haplotype assignment differs from truth, divided by the total number of covered variants; unlike SER, it counts each incorrect position independently rather than transitions between consecutive sites. c) The structural variant F1-score, computed by `truvari bench` [21], is the harmonic mean of precision and recall over variant calls matched between truth and test VCFs; the inferred VCF is restricted to sites carrying at least one alternate allele (removing homozygous-reference genotypes that do not constitute variant assertions) before matching against the full truth.

### 3.3 Computational performance

Among the four tools, `DipGenie` is the most computationally demanding: its geometric-mean runtime is 31 ×, 37 ×, and 45 × that of `VG`; 9.0 ×, 10.7 ×, and 12.0 × that of `Paragraph + Beagle`; and 10.4 ×, 11.8 ×, and 13.7 × that of `PanGenie + Beagle` at 1 ×, 2 ×, and total coverage respectively, though its peak memory footprint remains comparable to both genotyping pipelines (Figure 4). The slowdown increases with coverage because `DipGenie`’s runtime and memory scale with sequencing depth: deeper coverage produces more *k*-mers for the algorithm to evaluate, whereas the other three tools exhibit near-constant runtime and memory regardless of coverage. Even in the most expensive case (HG002, 12.41× depth, N50 = 251 bp; Supplementary Table A1), `DipGenie` completes in under 17 minutes with a peak RAM usage of 37 GB, well within the capacity of a single compute node.

**Fig. 4.**
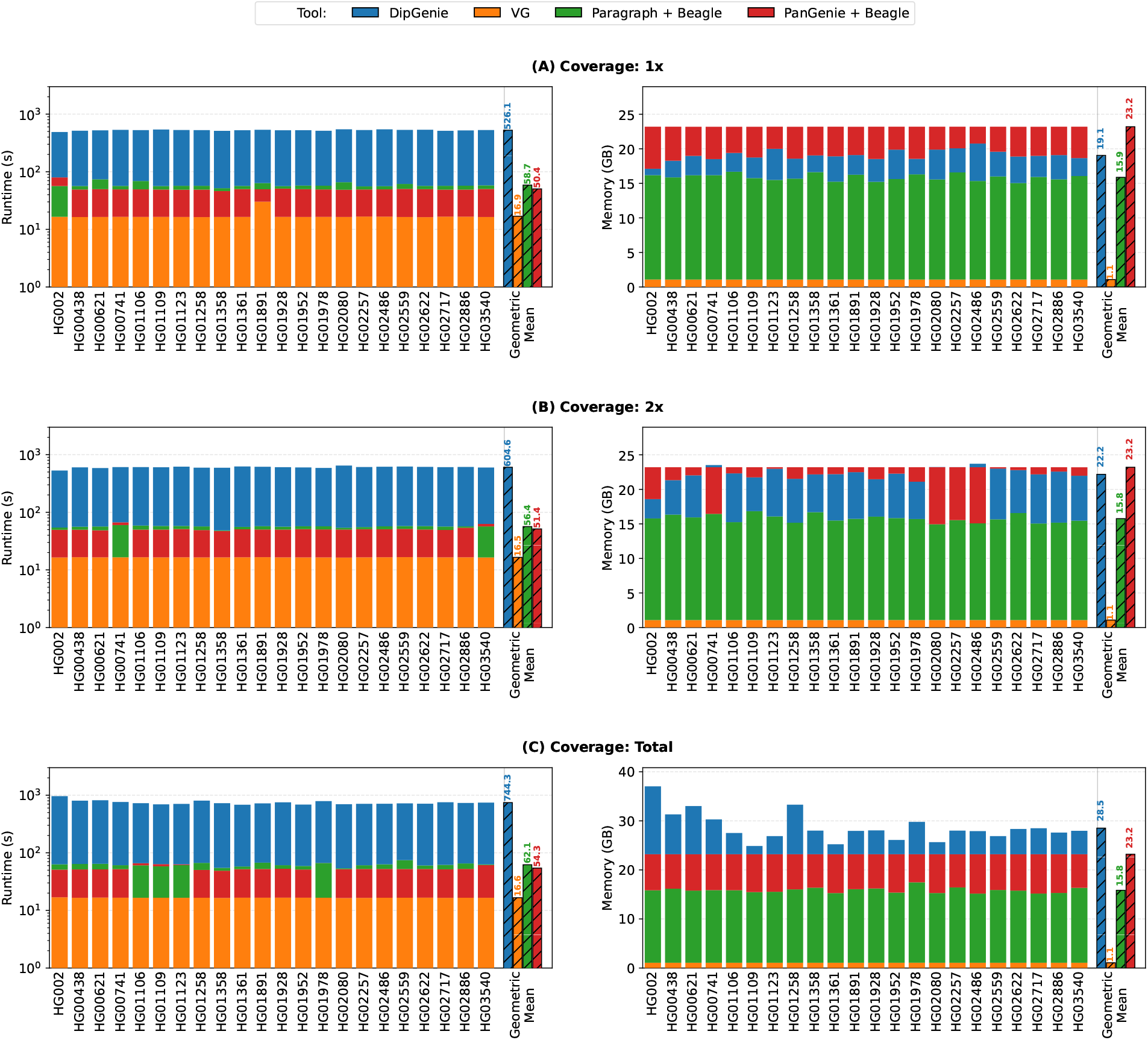
Runtime (left) and peak memory (right) for all four tools across 22 diploid samples at (A) 1*×*, (B) 2*×*, and (C) total coverage. Stacked bars show per-sample values; hatched bars to the right of the vertical separator indicate the geometric mean across all samples.

### 3.4 Structural variant calling

`DipGenie` leads in structural variant F1-score at every coverage level with GM of 0.504 (1×), 0.535 (2×), and 0.571 (total), exceeding `VG` (0.319, 0.351, 0.450), `PanGenie + Beagle` (0.318, 0.373, 0.470), and `Paragraph + Beagle` (0.275, 0.302, 0.379) by margins of 0.10–0.23 (Figure 6). All four tools produce evaluable SV calls for all 22 samples at every coverage level, so geometric means are computed over the full set. Precision drives this separation: `DipGenie` achieves GM precision of 0.597 at 1×, 0.611 at 2×, and 0.641 at total, compared to `VG` (0.274, 0.296, 0.371), `PanGenie + Beagle` (0.339, 0.402, 0.518), and `Paragraph + Beagle` (0.466, 0.523, 0.577). `VG` achieves the highest recall among all tools (0.385, 0.434, 0.583) but its low precision limits the resulting F1-score. `Paragraph + Beagle` shows precision second only to `DipGenie` but recall never exceeds 0.285, failing to recover over half the true variants. The SV counts explain this pattern (Supplementary Figure A1): against a ground-truth GM of 111.75 SVs per sample, `DipGenie` calls 139.74 (25.0% over-count), `PanGenie + Beagle` calls 87.57 (21.6% under-count), `VG` calls 73.85 (33.9% under-count), and `Paragraph + Beagle` calls 50.15 (55.1% under-count). `DipGenie`’s over-calling introduces some false positives but retains most true variants, whereas the three competing tools under-call to varying degrees, systematically missing true structural variants.

The F1-score differences above also vary with sequencing depth, quantifying each tool’s sensitivity to coverage (Figure 5). `DipGenie`’s F1-score improves by only 0.067 (0.504 to 0.571), the smallest gain among all tools, with precision rising 0.044 and recall 0.081, indicating that even at 1 × coverage its graph-based dynamic programming extracts sufficient haplotype signal for SV genotyping. `VG` shows the largest recall swing of 0.198 between 1 × and total, consistent with a method that requires dense read evidence to distinguish heterozygous from homozygous genotypes. `PanGenie + Beagle` is the most coverage-dependent tool overall, with precision increasing from 0.339 to 0.518 and recall from 0.302 to 0.433, an F1-score gain of 0.152 that still trails `DipGenie` at every level. `Paragraph + Beagle` improves precision by 0.111 (0.466 to 0.577) but recall remains lowest regardless of coverage. In practical terms, `DipGenie` at 1 × coverage (F1-score 0.504) already exceeds what `VG` and `PanGenie + Beagle` achieve at total coverage (0.450 and 0.470), reducing the sequencing investment required for reliable structural variant detection in the MHC.

**Fig. 5.**
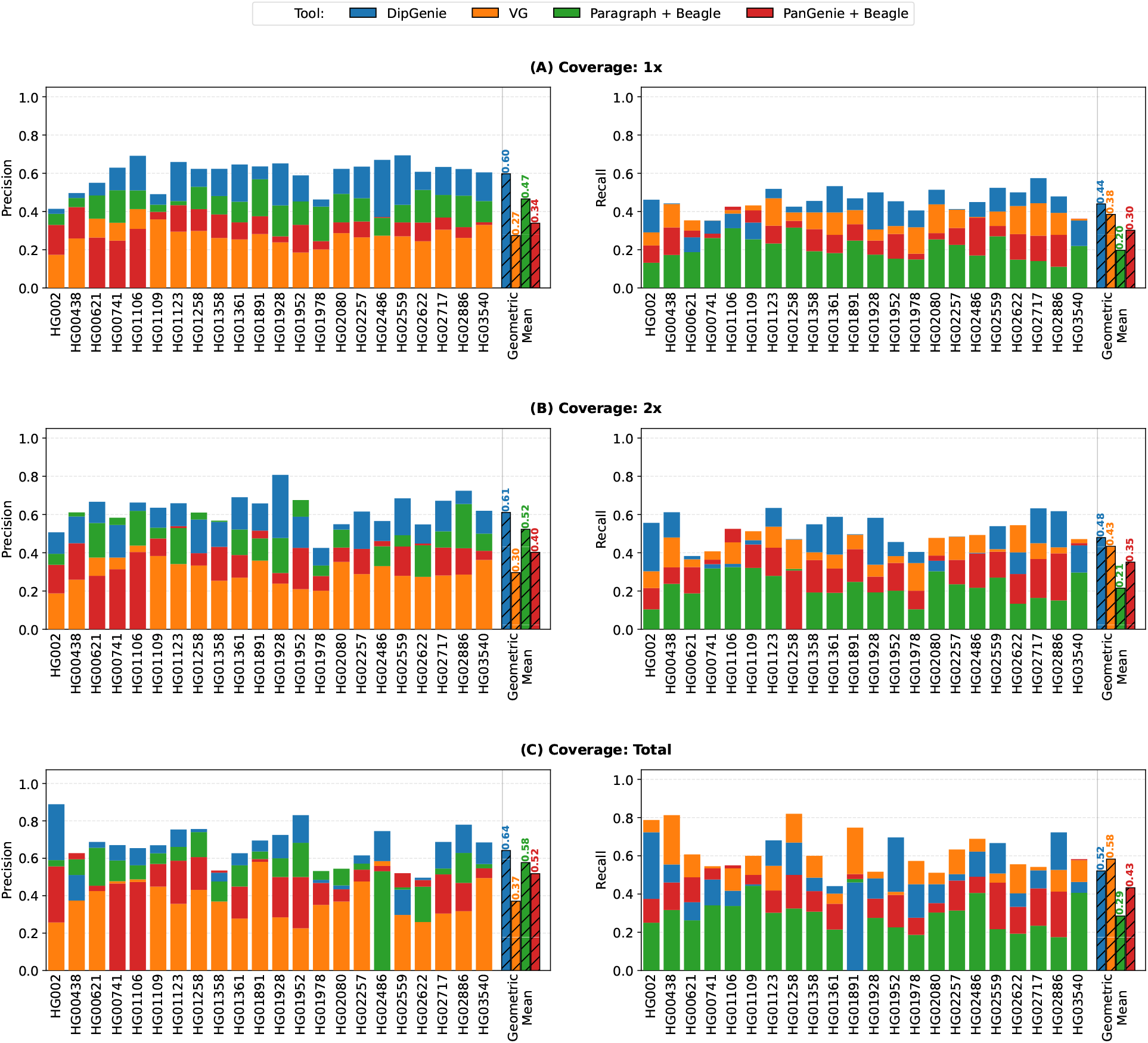
Precision (left) and recall (right) of structural variant calls for all four tools across 22 diploid samples at 1*×* (A), 2*×* (B), and total (C) coverage. Stacked bars show per-sample values; hatched bars to the right of the vertical separator indicate the geometric mean.

`DipGenie`’s balanced precision–recall profile traces to its recombination budget, which constrains inferred haplotypes to biologically plausible paths. At total coverage, the per-haplotype recombination counts converge toward the theoretically expected value of 9 (H1 GM = 8.68, H2 GM = 8.93; see Expected number of recombinations; Supplementary Figure A2). `VG` has no analogous constraint [52], and the panel-based pipelines impose no recombination structure at all.

### 3.5 Phasing quality

`DipGenie` achieves the lowest switch error rate at every coverage level. `VG` fails to produce heterozygous calls for 9 of 22 samples at 1 ×, 2 at 2 ×, and 4 at total coverage (it outputs homozygous genotypes when coverage is insufficient to distinguish haplotypes), and `Paragraph + Beagle` fails for 1 × sample at 1 (HG01258). To ensure a fair comparison, the figure reports geometric means over the intersection of samples for which all four tools produce evaluable phased output, reducing the common set to 12 at 1 ×, 20 at 2 ×, and 18 at total coverage. Over these common samples, `DipGenie`’s GM SER is 2.25% (1 ×), 1.44% (2 ×), and 0.86% (total), compared with `PanGenie + Beagle` at 6.44%, 5.66%, and 4.88%; `VG` at 5.44%, 6.92%, and 6.77%; and `Paragraph + Beagle` at 13.17%, 13.80%, and 11.35%. At total coverage, `DipGenie` is 5.7 × lower than `PanGenie + Beagle`, 7.9 × lower than `VG`, and 13.2 lower than `Paragraph + Beagle`. The advantage is already present at 1 ×, where `DipGenie`’s SER of 2.25% falls below the best competitor’s total-coverage value (`PanGenie + Beagle`, 4.88%), and it strengthens monotonically at 2 × (Figure 6). `DipGenie` infers phase jointly along entire haplotype paths, whereas `VG` phases independently within local graph blocks [52] and the genotyping pipelines delegate phasing to a separate statistical step that receives no graph-structural information [4].

**Fig. 6.**
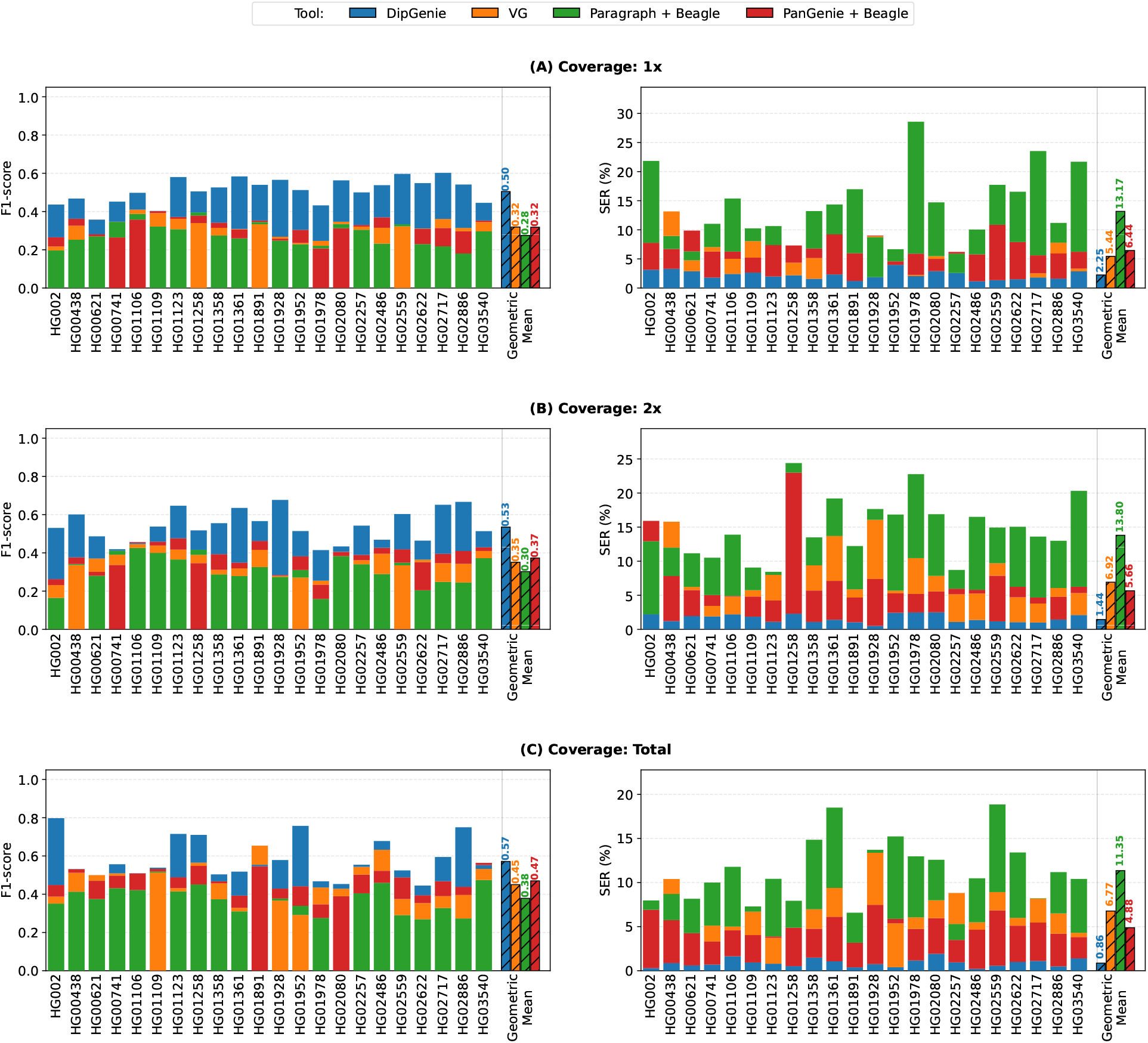
Structural variant F1-score (left; higher is better) and switch error rate (%, right; lower is better) across 22 diploid samples at (A) 1*×*, (B) 2*×*, and (C) total coverage. Stacked bars show per-sample values; hatched bars to the right of the vertical separator indicate the geometric mean over the common set of samples with evaluable output from all four tools (12 at 1*×*, 20 at 2*×*, 18 at total). `VG` produces zero heterozygous calls for a subset of samples (9 at 1*×*, 2 at 2*×*, 4 at total) and is therefore absent from those per-sample bars; `Paragraph + Beagle` fails for 1 sample at 1*×*.

Mismatch rates measure per-position allele-to-haplotype assignment accuracy within phased blocks (Supplementary Figure A3). Over the same common sample sets, `DipGenie`’s GM mismatch rate at 1 × is 37.00%, positioned between `VG` (31.15%) and `PanGenie + Beagle` (39.56%), with `Paragraph + Beagle` at 32.35%. As coverage increases, `VG`, `Paragraph + Beagle`, and `PanGenie + Beagle` all worsen: their GM mismatch rates rise to 35.09%, 40.42%, and 45.38% at total coverage. `DipGenie`’s rate instead falls monotonically from 37.00% to 32.36% (2 ×) and 21.66% (total), a 1.71 × self-improvement from 1 × at total coverage. At full depth `DipGenie` is 1.62 × more accurate than `VG`, 1.87 × than `Paragraph + Beagle`, and 2.10 × than `PanGenie + Beagle`, making it the only method tested that consistently converts additional sequencing into lower mismatch rates.

### 3.6 Approximation quality

A natural question is how close our heuristic comes to the true optimum. We address this by computing an *a posteriori* approximation ratio that upper-bounds obj(*P* ^∗^)*/* obj(*P*), where *P* ^∗^ is an optimal path pair and *P* is the heuristic solution; a value of 1 would certify optimality. The certificate

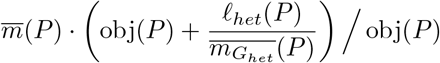

is guaranteed to be an upper bound by Theorem 1, and all quantities are computed directly from *P* . Across our leave-one-out experiments, the GM ratios are 6.45, 5.52, and 4.99 at 1 ×, 2 ×, and total coverage respectively (Figure 7). The ratio decreases with coverage because deeper sequencing provides more informative *k*-mer counts, allowing the algorithm to approach the optimum more closely. The highest individual ratios appear at 1 × (e.g. HG002, 8.17), where sparse *k*-mer evidence increases the color multiplicity term 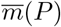 and loosens the bound; reassuringly, no corresponding degradation appears in SER or F1-score, indicating that the bound overestimates the true gap.

**Fig. 7.**
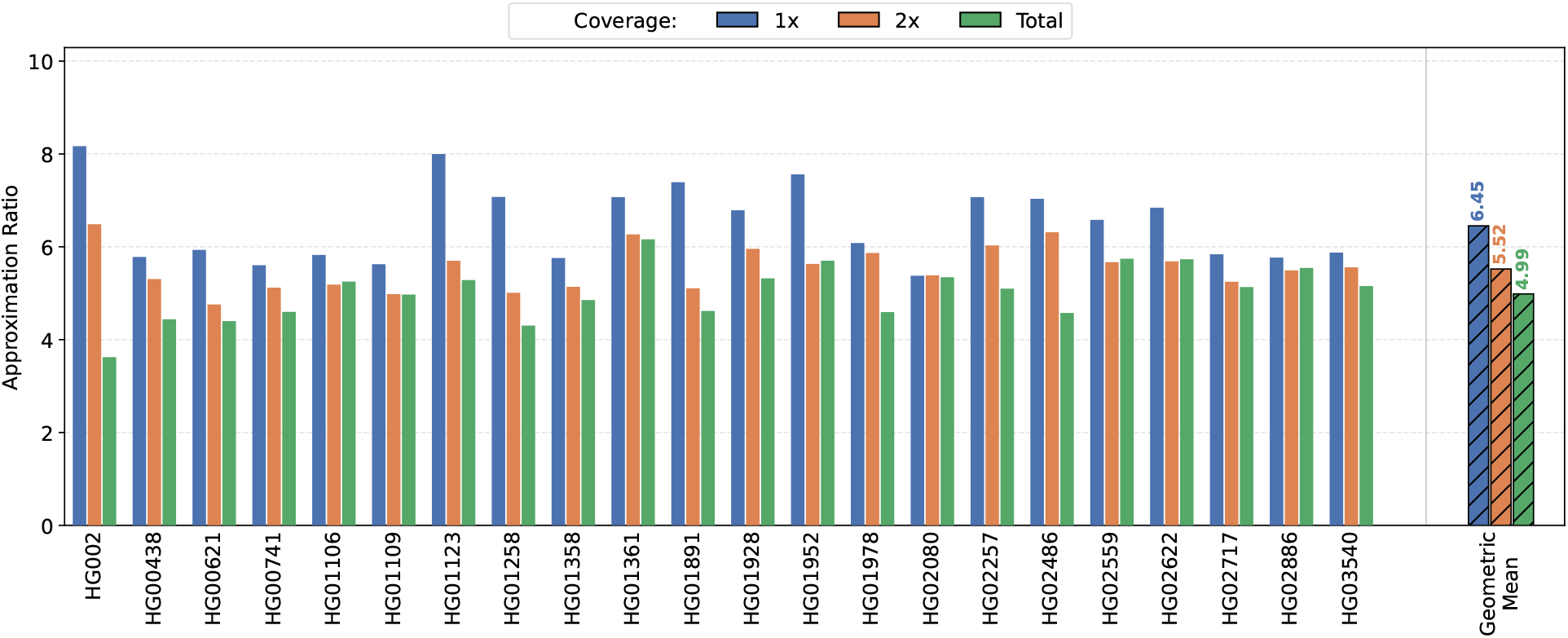
*A posteriori* approximation ratio (upper bound on obj(*P* ^∗^)*/* obj(*P*)) across 22 leave-one-out experiments at 1*×*, 2*×*, and total coverage. Grouped bars show per-sample values; hatched bars to the right of the vertical separator indicate the geometric mean at each coverage level. Lower values indicate solutions closer to the theoretical optimum; the ratio improves monotonically with coverage.

## 4 Discussion

Genotyping is a fundamental task in computational biology, supporting downstream applications in genomics and genetics [38, 39]. Pangenome graphs offer a compact representation of population-level haplotype diversity, capturing SNPs, indels, and structural variants in a single structure, and have become a standard platform for variant analysis [1, 2, 13, 18, 19, 26–28, 37, 38, 42, 51, 55, 56]. Population-level statistical methods perform imputation and phasing jointly but operate on linear haplotype reference panels without graph structure [4, 15, 31, 40, 48]. Graph-based methods genotype variants using the pangenome but achieve only block-wise phasing within local graph regions [10, 18, 52]. Neither approach jointly optimizes genotyping and phasing along global paths through the pangenome graph. This limitation is especially acute in the MHC, where dense polymorphism and extended linkage disequilibrium demand globally coherent haplotype inference [16]. We addressed this gap by formulating diploid genome inference as a combinatorial optimization problem on pangenome graphs and developing a tool that solves this problem via sequential dynamic programming with a biologically motivated recombination budget and provable *a posteriori* approximation guarantees. Across 22 diploid leave-one-out experiments in the MHC, our method achieves GM switch error rates up to 13.2 × lower than the best competitor, leads in structural variant F1-score at every coverage level (0.504–0.571 vs 0.275–0.470), and is the only method tested whose mismatch rate improves monotonically with sequencing depth. These gains come at higher runtime and memory, which remains practical on a single compute node but motivates the directions below.

Whole-genome pangenome graphs constructed from multiple assemblies typically contain cycles, primarily due to large inversions [23, 25, 28, 37]. Our current dynamic programming relies on the topological-sort traversal of a DAG; extending it to cyclic graphs requires strict biologically inspired constraints that limit cycle traversal to plausible copy numbers. Our current experiments use `DipGenie` in the MHC region as a proof of concept to demonstrate a new approach to diploid genome inference. In the MHC graph, every haplotype spans an end-to-end alignment from source to sink, so the algorithm can trace complete paths. In whole-genome graphs, however, end-to-end alignments are often missing because long inversions, segmental duplications, and other complex rearrangements break the contiguity of individual haplotype paths through the graph. `DipGenie`’s algorithm currently cannot handle such broken paths, which prevents direct application to whole-genome graphs. In future work, we aim to extend `DipGenie` to operate on fragmented haplotype paths and cyclic structures, enabling it to scale to whole-genome pangenome graphs while preserving the phasing accuracy demonstrated here.

## Appendix

### A1 ILP formulation

The ILP formulation for Diploid path inference can be obtained by modifying the ILP formulation given by Chandra et al. [8]. In particular, we must change the objective function and add a single constraint. The objective function should be changed to ∑_*r*∈S_ *z*_*r*_ (our notation for the variable *z*_*r*_ used below is kept consistent with that previous work). The constraint ∑_(*u,v*)∈*E*_ *w*(*u, v*) · *x*_*u,v*_ ≤ *R* should also be added, where *w*(*u, v*) is the weight for edge (*u, v*).

Let *G*_*E*_ = (*V*_*E*_, *E*_*E*_) be the expanded graph, let *w*(*u, v*) denote the weight of an edge (*u, v*) ∈ *E*_*E*_, and let 𝒮= 𝒮_*het*_ ∪ 𝒮_*hom*_. For each target string *s* recall that hit(*s*) denotes the set of all paths corresponding to *s* in *G*_*E*_. For *p* ∈ hit(*s*), we write |*p*| for the number of edges in *p*. We write 𝒩^+^(*u*) and 𝒩^−^(*u*) for the outgoing and incoming neighbors of *u*, respectively.

An ILP formulation is:

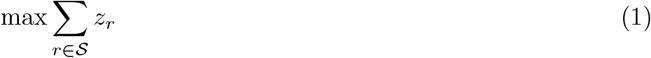

subject to the variable domains

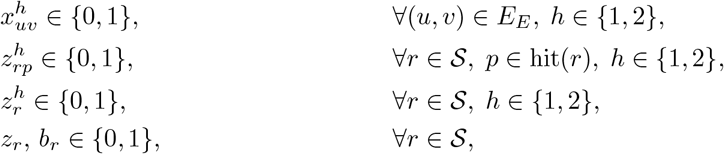

and constraints:

#### Path constraints

For each *h* ∈ {1, 2}, the 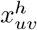 encode an *s* → *t* path: ∀*u* ∈ *V*_*E*_, *h* ∈ {1, 2}

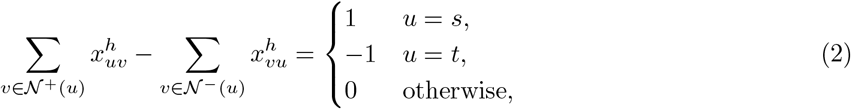

#### Recombination budget

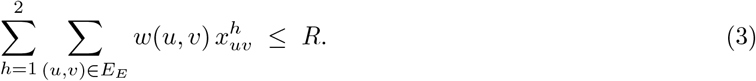

#### String–path matching

For every *r* ∈ 𝒮, *p* ∈ hit(*r*), *h* ∈ {1, 2}:

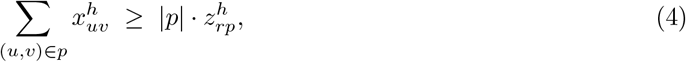

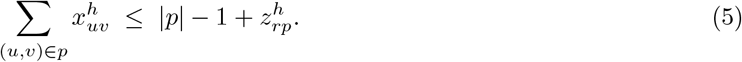

Together these enforce:

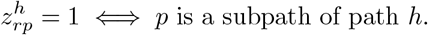

#### String-level activation

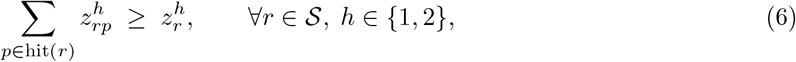

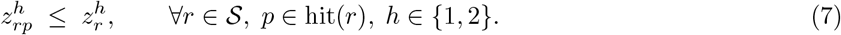

Thus 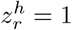 iff some *p* ∈ hit(*r*) appears on path *h*.

#### Homozygous scoring

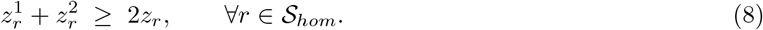

#### Heterozygous scoring

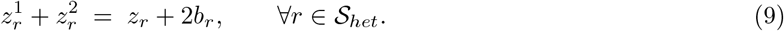

In the objective (1), each *z*_*r*_ = 1 contributes one unit of score. Constraints (2)–(3) ensure the solution paths form feasible *s* → *t* paths under the shared recombination budget. Constraints (4)–(5) encode path containment. Constraints (6)–(7) activate 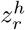 exactly when *r* appears on haplotype *h*. Finally, (8) enforces that homozygous strings count only if they appear on both haplotypes, while (9) enforces that heterozygous strings count only if they appear on exactly one haplotype.

### A2 Proof of Theorem 1

Let *X* = (*X*_1_, *X*_2_) denote any pair of paths. The DP score on *X* can be interpreted as considering colors from different levels as being distinct, while maintaining their *hom/het* classification within a given level. We define the following quantities.

#### Homozygous quantities

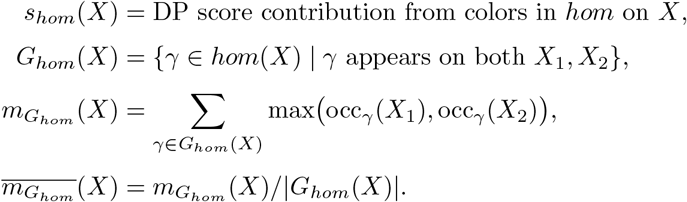

#### Heterozygous quantities

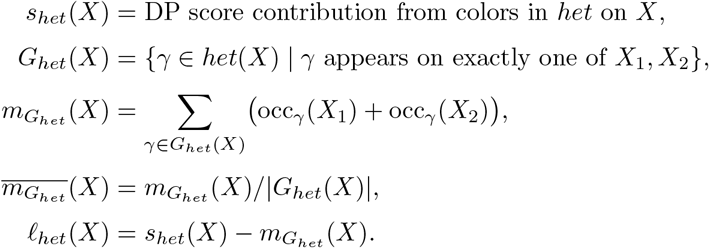

##### Theorem 1.

*Let P be a solution path pair obtained by our algorithm for Problem 2, and let P* ^∗^ *be an optimal solution. Let* 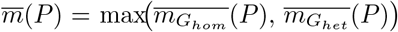 *be the maximum average multiplicity of contributing homozygous or heterozygous colors on P, and let ℓ*_*het*_(*P*) *denote the heterozygous score loss. Then*

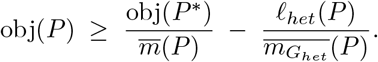

*Proof*. For homozygous colors,

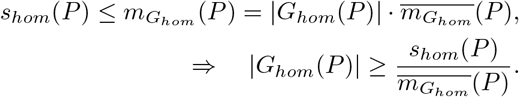

For heterozygous colors,

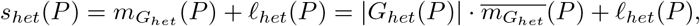

hence

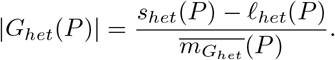

Combining,

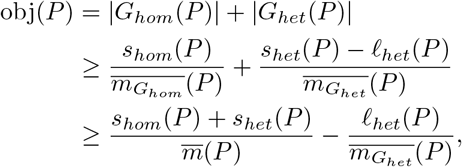

where 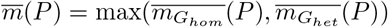.

Since the DP algorithm maximizes *s*_*hom*_ + *s*_*het*_,

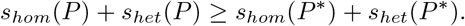

Each color *γ* counted in |*G*_*het*_(*P* ^∗^)| contributes at least 1 to *s*_*het*_(*P* ^∗^), since there must exist some level where *γ* appears on only *v*_1_ or *v*_2_. This implies |*G*_*het*_(*P* ^∗^)| ≤ *s*_*het*_(*P* ^∗^). Each color *γ* counted (potentially multiple times) in *s*_*hom*_(*P* ^∗^) contributes at most 1 to |*G*_*hom*_(*P* ^∗^)|, since repeated occurrences of *γ* are not counted in |*G*_*hom*_(*P* ^∗^)|. This implies *s*_*hom*_(*P* ^∗^) ≥ |*G*_*hom*_(*P* ^∗^)|. These together imply

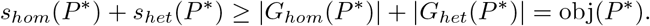

Therefore,

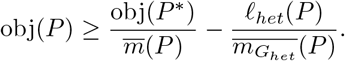

We see that if *ℓ*_*het*_(*P*) = 0, that is, the heterozygous loss is 0 and 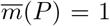, that is, all colors appear with multiplicity 1, the approximate solution is optimal.

### A3 Expected number of recombinations

We estimate the expected number of recombinations using the Li and Stephens (2003) model [36], which represents haplotype copying as a Hidden Markov Model (HMM). At each site *j*, the hidden state *Z*_*j*_ ∈ {1, …, |ℋ|} indicates the donor haplotype being copied, with transition probability

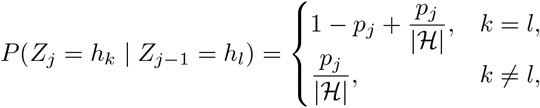

where

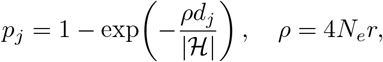

and *d*_*j*_ is the physical distance between adjacent sites.

A recombination corresponds to a change in copying donor haplotype. Based on the above, the probability of recombination is:

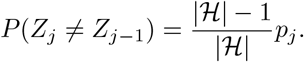

Assuming constant per-base distance *d*_*j*_ = 1 bp and uniform *p*_*j*_ = *p*, the expected number of recombinations across all sites of a reference of length *L* is

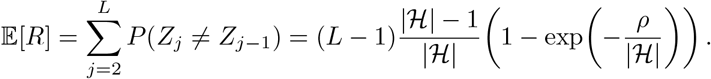

Substituting *ρ* = 4*N*_*e*_*r* yields

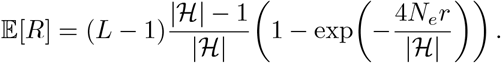

#### A3.1 Parameter values used in experiments

– Reference length: *L* = 4,920,303 bp.
– Number of reference haplotypes: |ℋ| = 47.
– Effective population size: *N*_*e*_ = 5 × 10^3^ [44].
– Recombination rate: *r* = 0.46 cM/Mbp = 4.6 × 10^−9^ Morgans/bp [53].

These parameter values make 𝔼[*R*] ≈ 9.

**Table A1.**
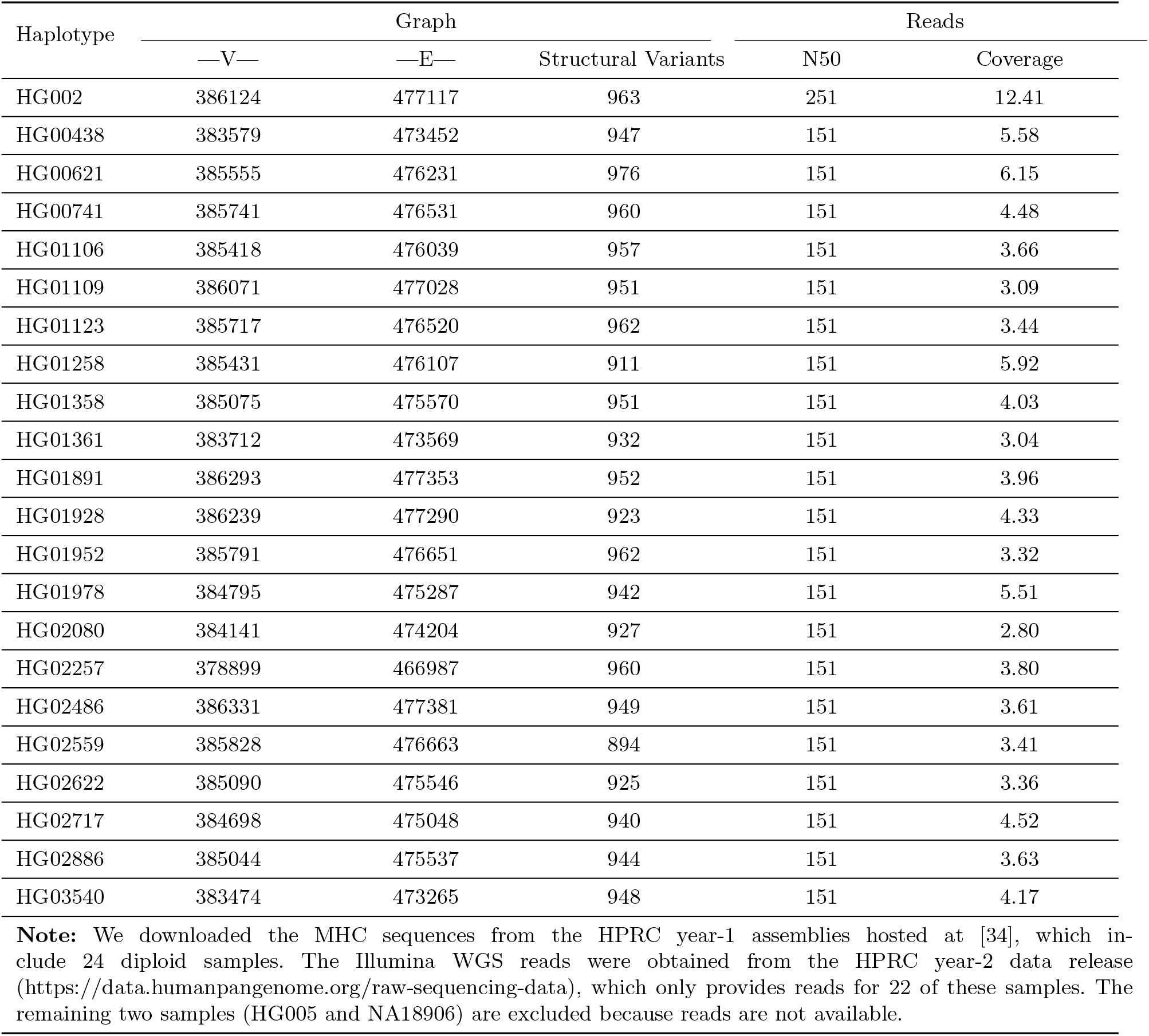
Statistics for the graphs and read sets used in the leave-one-out experiments. For each experiment, we construct a graph by excluding the haplotype sample being evaluated; thus, every graph contains the remaining 23 diploid samples plus CHM13 as the reference, for a total of 47 haplotypes. We report the corresponding graph statistics. For the read sets, we only report the N50 and the total sequencing coverage.

**Fig. A1.**
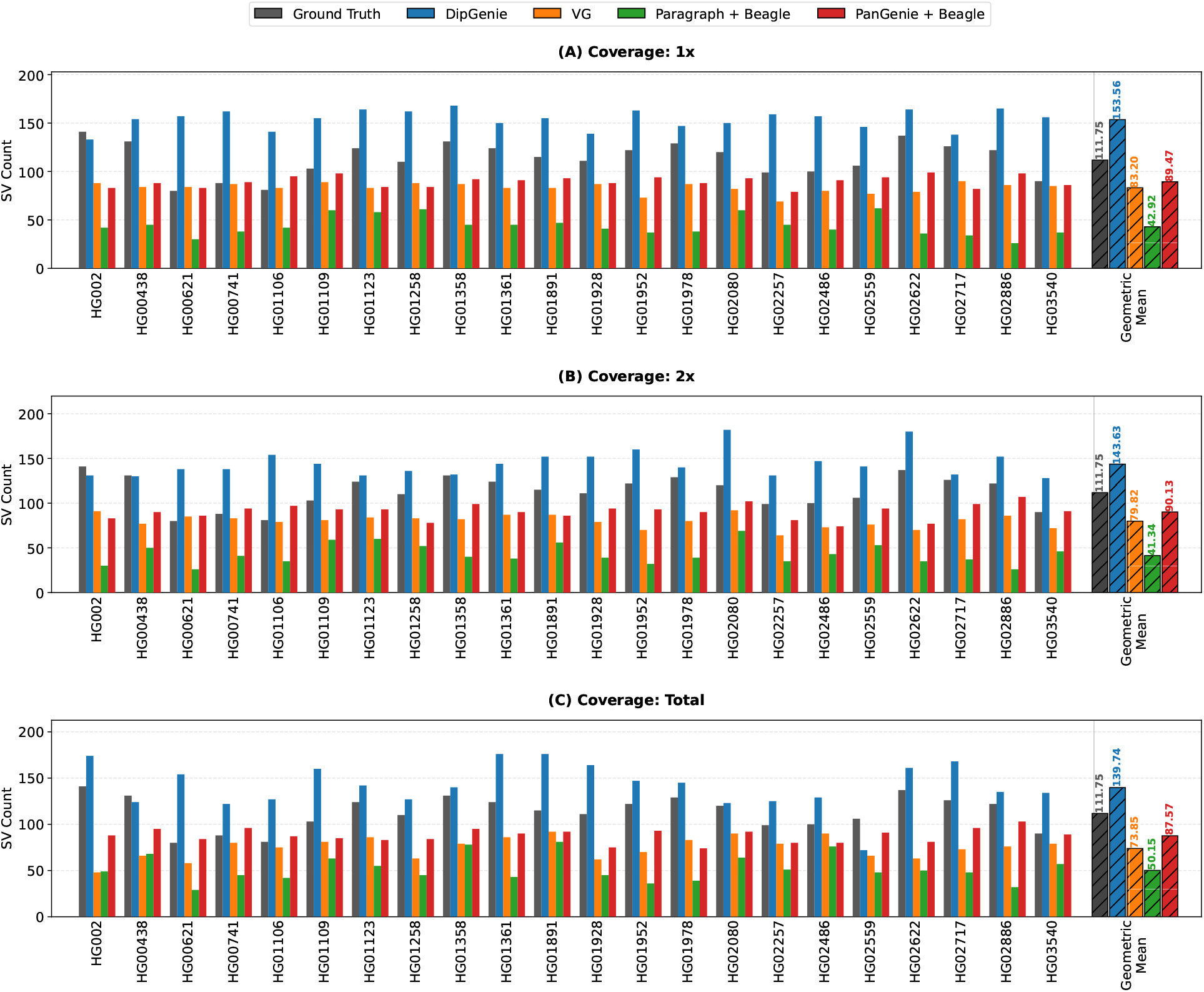
The figure illustrates the count of structural variants (SVs) from the leave-one-out experiment conducted for various haplotype samples and sequencing coverages (1*×*, 2*×*, and Total). The hatched Geometric Mean (GM) bar summarizes SV counts across all 22 samples. Against a ground truth GM of 111.75, DipGenie is the only tool that consistently exceeds the ground truth (125.0%–137.4% recovery), suggesting higher sensitivity at the cost of potential false positives. All other tools under-count SVs. PanGenie + Beagle recovers 78.4%–80.7% of the ground truth, VG recovers 66.1%–74.4%, and Paragraph + Beagle captures only 37.0%–44.9%. DipGenie, VG, and PanGenie + Beagle show monotonically decreasing SV counts from 1x to Total coverage, while Paragraph + Beagle decreases from 1*×* (GM: 42.92) to 2*×* (GM: 41.34) but increases at Total coverage (GM: 50.15).

**Fig. A2.**
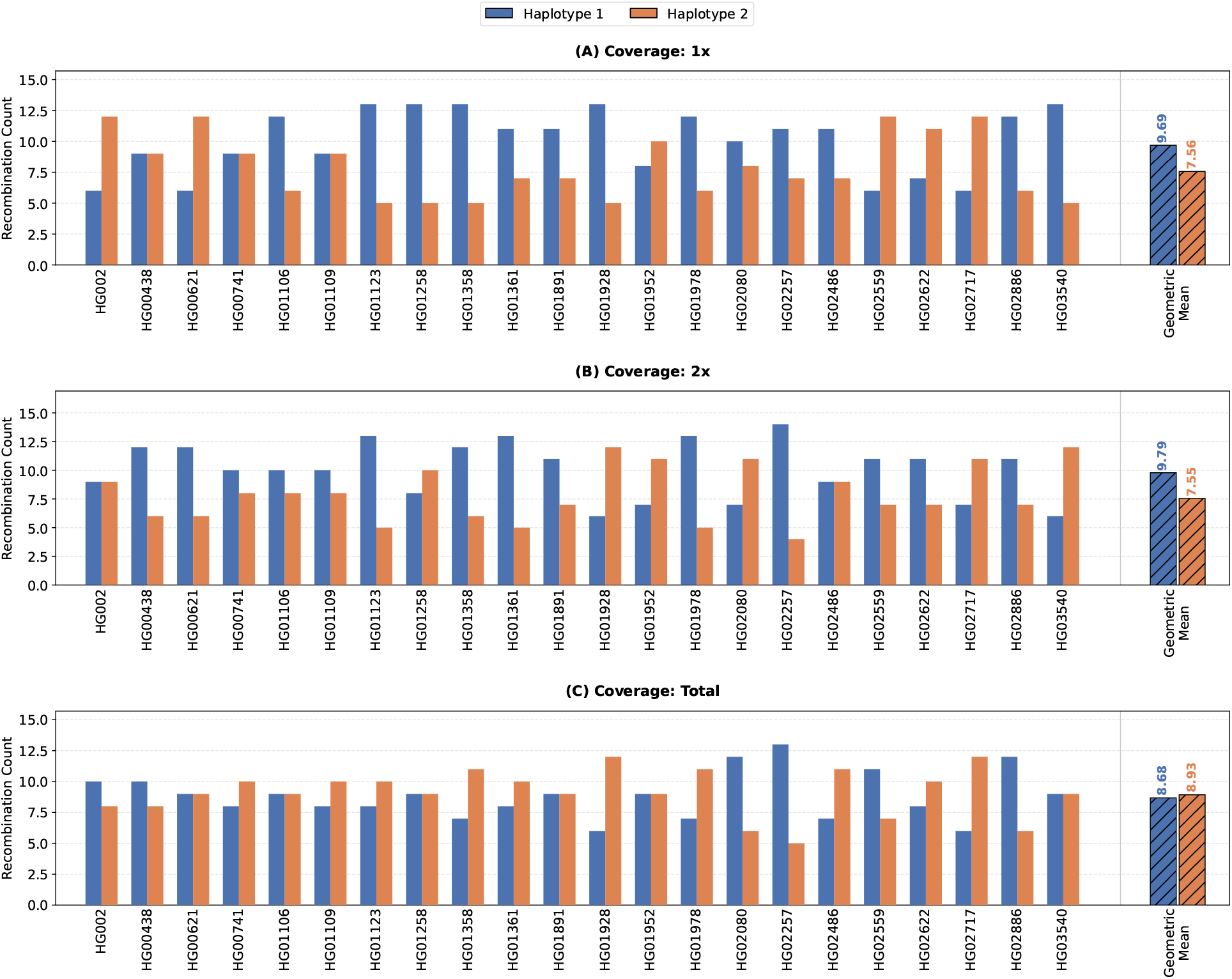
The figure shows recombination counts for the two diploid haplotypes, Haplotype 1 (H1) and Haplotype 2 (H2), inferred by DipGenie across different read coverages (1*×*, 2*×*, and Total). The total recombination count per sample is constrained to H1 + H2 ≤ 18 as an input parameter. The theoretically expected number of recombinations is 9 per haplotype A3. At 1*×* and 2*×* coverage, the per-haplotype Geometric Mean (GM) values are imbalanced (1*×*: H1 GM = 9.69, H2 GM = 7.56; 2*×*: H1 GM = 9.79, H2 GM = 7.55), whereas at Total coverage the GM values converge toward the expected value of 9 (H1 GM = 8.68, H2 GM = 8.93), indicating that higher coverage enables more accurate partitioning of recombinations between the two haplotypes.

**Fig. A3.**
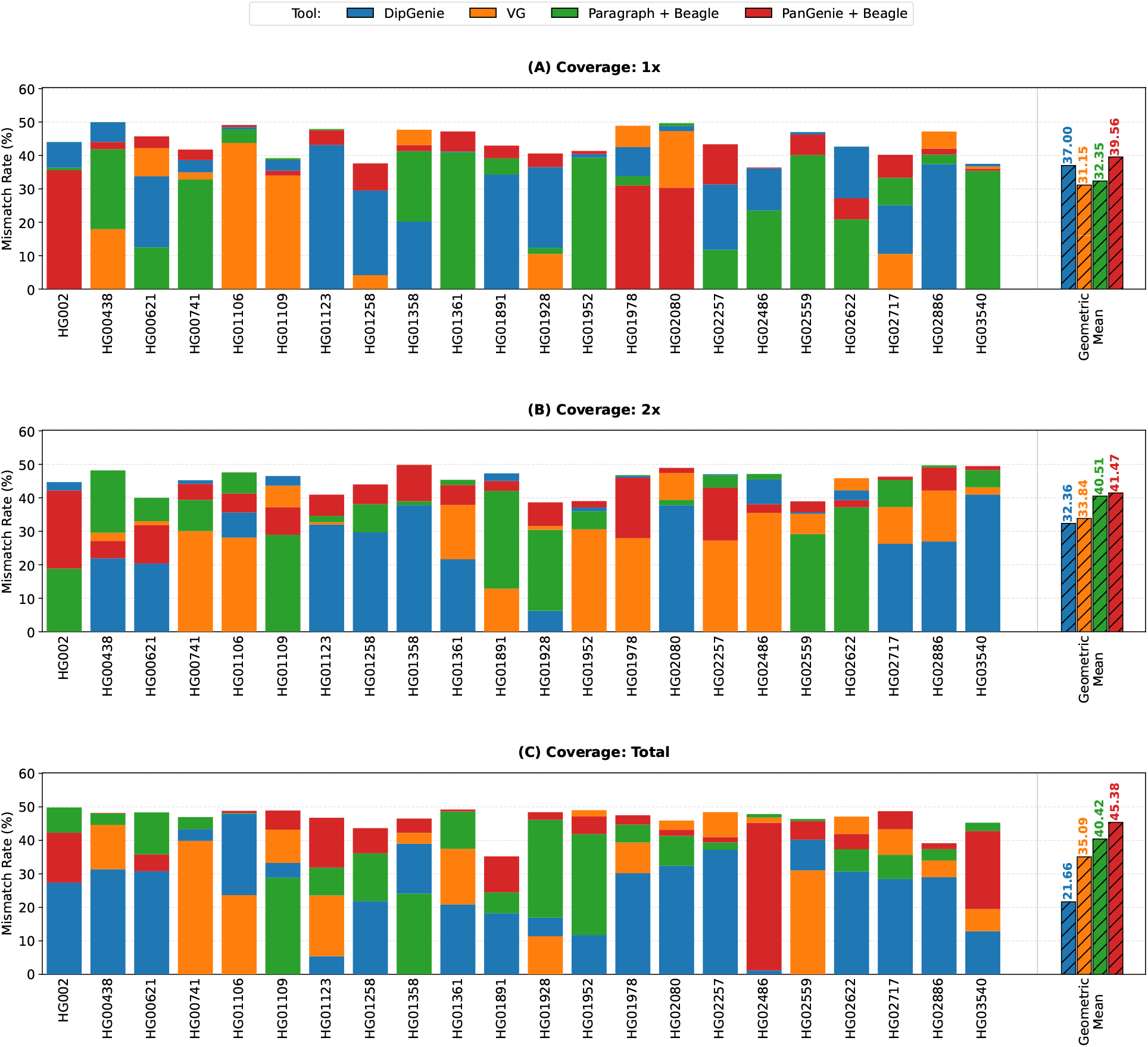
Mismatch rate (%), defined as the percentage Hamming distance between inferred and true haplotype assignments at heterozygous positions [41], for four tools across 22 diploid samples at 1*×*, 2*×*, and Total coverages. Lower values indicate better accuracy. The hatched bar shows the Geometric Mean (GM) across samples. At 1*×* coverage, VG has the lowest GM (26.69%), followed by Paragraph + Beagle (31.71%), DipGenie (37.70%), and PanGenie + Beagle (39.78%). As coverage increases, DipGenie improves monotonically (37.70% → 32.71% → 22.08%; 1.71*×* reduction), while the other tools worsen: VG 26.69% → 35.09%, Paragraph + Beagle 31.71% → 40.01%, PanGenie + Beagle 39.78% → 44.16%. At Total coverage, DipGenie achieves the lowest GM (22.08%), which is 1.59*×*, 1.81*×*, and 2.00*×* lower than VG, Paragraph + Beagle, and PanGenie + Beagle, respectively. DipGenie is the only tool that leverages additional sequencing depth to improve haplotype accuracy.

